# BBATProt : A Framework Predicting Biological Function with Enhanced Feature Extraction via Explainable Deep Learning

**DOI:** 10.1101/2024.10.16.618767

**Authors:** Youqing Wang, Xukai Ye, Yue Feng, Haoqian Wang, Xiaofan Lin, Xin Ma, Yifei Zhang

## Abstract

Accurately predicting the functions of peptides and proteins from their amino acid sequences is essential for understanding life processes and advancing biomolecule engineering. Due to the time-consuming and resource-intensive nature of experimental procedures, computational approaches, especially those based on machine learning frameworks, have garnered significant interest. However, many existing machine learning tools are limited to specific tasks and lack adaptability across different predictions. Here we propose a versatile framework BBATProt for the prediction of various protein and peptide functions. BBATProt employs transfer learning with a pre-trained Bidirectional Encoder Representations from Transformers (BERT) model, to effectively capture high-dimensional features from amino acid sequences. The whole custom-designed network, integrating Bidirectional Long Short-Term Memory (Bi-LSTM) and Temporal Convolutional Networks (TCN), can align with the spatial characteristics of proteins. It combines local and global feature extraction through attention mechanisms for precise functional prediction. This approach ensures that key features are adaptively extracted and balanced across diverse tasks. Comprehensive evaluations show BBATProt outperforms state-of-the-art models in predicting functions like hydrolytic catalysis, activity of peptides, and post-translational modification sites. Visualizations of feature evolution and refinement via attention mechanisms validate the framework’s interpretability, providing transparency into the evolutional process and offering deeper insights into function prediction.

## Introduction

Protein and peptide, the biomacromolecules composed of amino acid strings, execute a diverse array of essential biological functions in living organisms. Unraveling how these strings encode the structural and topological characteristics that determine their bioactivities is the core of biology. Previous studies, relying heavily on cumbersome wet experiments such as protein crystallography and biochemical assays, have advanced slowly (1). For instance, of the more than 100 million sequences recorded in the UniProt database, less approximately 0.5% have undergone manual annotation (UniProtKB/Swiss-Prot) (2). Despite AlphaFold’s accomplishment in demonstrating that amino acid sequences encompass all necessary information for folding, accurately predicting their functions purely from these sequences remains difficult (3).

The difficulty in directly mapping a sequence of amino acids to its potential function arises from the complexity and diversity of functions (4). A particular biological function relies not only on the specific composition or configuration of key residues but is also influenced in varying degrees by other residues in the vicinity or even far away (5). For instance, the catalytic activity of an enzyme is mostly controlled by the residues in the catalytic pocket but may also be regulated by remote residues. The reactivity of a post-translational modification (PTM) site of a protein depends on the chemical environment of the target functional group, as well as the local landscape. Since peptides are considerably smaller than proteins, factors such as their composition, sequence patterns, and overall shapes all contribute to biological functions to different extents. Therefore, accurately predicting biological functions based on sequences requires an adaptive framework capable of automatically balancing multiple-level features of chemical and structural characteristics (6).

Traditional shallow machine learning methods, including K-Nearest Neighbors (KNN), Random Forest (RF), support vector machine (SVM) and so on (7; 8; 9; 10), have been commonly utilized for predicting specific functions of peptides and proteins. However, these methods suffer from limited generalization capabilities due to their dependence on high-quality manually annotated features, which restrict their ability to capture implicit and high-dimensional information within sequences, ultimately hindering their adaptability to diverse datasets and novel contexts. In contrast, deep neural network approaches have emerged to effectively map the complex, high-dimensional, and nonlinear relationships between sequences and their biological functions. These methods include Convolutional Neural Networks (CNN), Bidirectional Long Short-Term Memory (Bi-LSTM), Temporal Convolutional Networks (TCN), and advanced Bidirectional Encoder Representations from Transformers (BERT) (11; 12; 13; 14; 15). Notably, transfer learning techniques, such as BERT, which excel in leveraging contextual understanding from one domain to another, have proven highly effective in adapting to protein sequence analysis, enabling the model to capture intricate biological patterns with minimal task-specific data. Rives et al. used 250 million protein sequences to pretrain a protein language model by considering amino acid sequences as sentences (16), and demonstrated BERT’s efficacy in capturing high-dimensional structural and functional information in protein sequences. Building on this architecture, models such as ProtBERT, ESM-1b, and BERT-Protein have been developed. These models extract high-dimensional abstract characteristics from protein sequences by using large-scale unsupervised learning and transfer learning methodologies (17; 18; 19).

Given the complexities inherent in protein and peptide function prediction, the development of a novel framework capable of extracting multi-level features that capture both local and global sequence information is imperative. In this study, we present BBATProt, a feature-enhanced network framework that utilizes BERT for effective dynamic word embedding extraction from amino acid sequences. The framework was custom-designed to incorporate a temporal data-based architecture that aligns with the interpretability of protein spatial structural relationships, thereby significantly enhancing the precision of function prediction. The workflow of BBATProt is illustrated in Fig. 1. To address the intricate interrelationships among high-dimensional features within sequences, BBATProt employs BERT for feature extraction, which adeptly captures contextual information inherent in the amino acid sequences. The incorporation of BERT facilitates the dynamic representation of sequences, allowing the model to adaptively learn from a vast array of protein sequences without the need for extensive prior knowledge of protein structures or functional domains. BBATProt’s design emphasizes multi-level feature extraction, which is crucial for understanding the diverse functionalities of proteins and peptides. The framework utilizes a targeted combination of CNN, Bi-LSTM, and TCN is proposed to effectively encapsulate the complex information embedded in the encoded features. This methodology proficiently manages long-range dependencies and high-dimensional abstract features, optimizing the extraction of correlations among features derived from temporal data while minimizing computational overhead. Furthermore, the self-attention mechanism optimally leverages interdependencies among features, reflecting the nonlinear significance of different sequence positions for functional realization. To assess the algorithm’s effectiveness, extensive evaluations across five independent datasets—including one hydrolytic enzyme dataset, two peptide datasets, and two post-translational modification (PTM) site datasets—reveal that BBATProt excels in accuracy, robustness, and generalization compared to current state-of-the-art (SOTA) models. Additioinaly to further validate the interpretability of the BBATProt architecture, t-distributed stochastic neighbor embedding (t-SNE) is used to illustrate the layer-by-layer evolution of features, while the refinement achieved through the attention mechanism is demonstrated separately. This study, grounded in dynamic word embeddings and neural networks, effectively identifies key functional features and showcases broad applicability and stability in practical scenarios, thus laying a solid foundation for future research in protein and peptide function prediction.

**Fig. 1.**
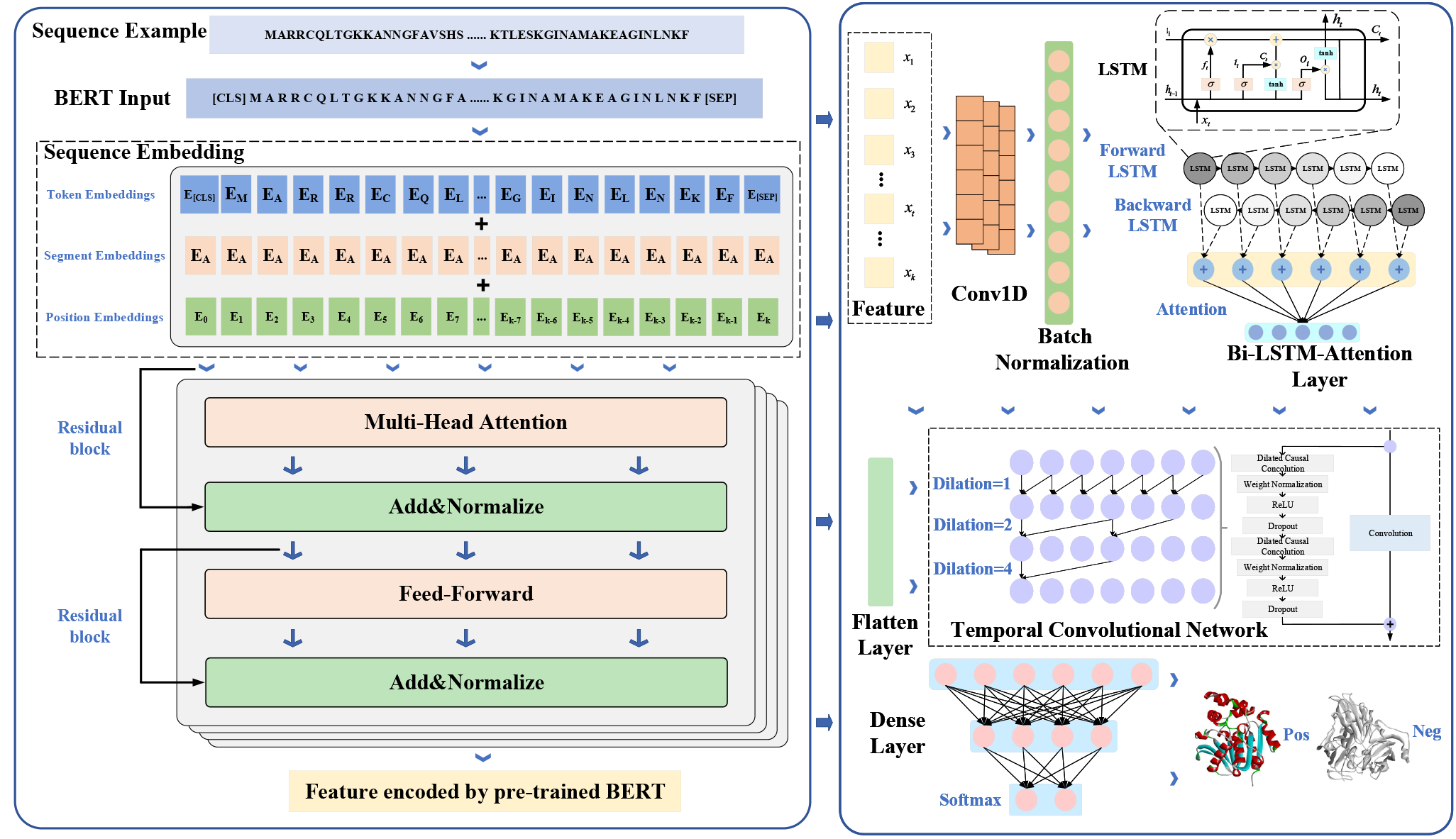
Overall framework of BBATProt

## Materials and Methods

### Dataset Construction

Due to the lack of a benchmark dataset for validating the model’s general versatility, Due to the lack of a benchmark dataset for validating the prediction model’s general versatility, a comprehensive series of comparative experiments was carried out across multiple benchmark datasets to illustrate the effectiveness of BBATProt. This study seeks to conduct a comprehensive assessment of the algorithm’s adaptability and robustness in a range of biological contexts. The framework was strategically designed to accommodate a range of functional prediction needs, encompassing instances such as antimicrobial peptides (AMP), dipeptidyl peptidase-IV (DPP-IV) inhibitory peptides, carboxylesterase hydrolysis prediction, and post-translational modification (PTM) site prediction (20; 21; 22; 23). To minimize redundancy within these datasets, a unique two-step operation deploying CD-HIT was executed at both protein and fragment levels (24). Table 1 presents the specific sources and detailed information of the datasets.

**Table 1.** Information on five peptide and protein datasets.

| Dataset | Training datasets |  | Independent datasets |  |
| --- | --- | --- | --- | --- |
|  | Pos | Neg | Pos | Neg |
| Hydrolytic enzyme<sup>(25)</sup> | 3132 | 3132 | 348 | 348 |
| Antimicrobial peptide<sup>(26)</sup> | 5536 | 5536 | 1536 | 1536 |
| Inhibitory peptide<sup>(27)</sup> | 532 | 532 | 133 | 133 |
| Lysine crotonylation<sup>(28)</sup> | 6975 | 6975 | 2989 | 2989 |
| Lysine glycation<sup>(29)</sup> | 3573 | 3573 | 200 | 200 |

**Table 2.** Comparative Representational Ability of BERT Encoding Features Across Different Sizes (%).

| Pre-trained model | ACC | MCC | SEN | SPE | PRE | FSc | AUROC |
| --- | --- | --- | --- | --- | --- | --- | --- |
| BERT-Tiny | 92.74 | 85.97 | 90.00 | **95.49** | **95.44** | 92.36 | 92.9 |
| BERT-Mini | 93.20 | 86.60 | 92.84 | 93.56 | 93.70 | 93.17 | 92.7 |
| BERT-Small | **94.21** | **88.54** | **93.48** | 94.94 | 94.96 | **94.15** | **94.21** |
| BERT-Medium | 93.07 | 86.31 | 92.24 | 93.91 | 93.95 | 93.01 | 93.10 |
| BERT-Base | 89.55 | 79.71 | 89.11 | 90.00 | 90.77 | 89.62 | 90.20 |
| BERT-Protein | 86.94 | 74.67 | 87.33 | 86.55 | 87.63 | 87.02 | 87.20 |

Note: The optimal performance for each metric is indicated in bold.

In the experiment, a 10-fold cross-validation approach was employed to evaluate the algorithm’s robustness and reliability. This methodology entails partitioning the dataset into ten distinct, non-overlapping subsets. During each iteration, training is conducted using nine subsets, while the remaining subset is reserved for independent testing. This iterative process facilitates a thorough assessment of the algorithm’s performance. The adoption of this cross-validation method helps minimize bias in the selection of training and test sets, thereby improving the scientific objectivity of the experimental results and ensuring replicability.

### BERT feature extraction

Pre-trained natural language processing models have found extensive application across various fields (30; 31). Transfer learning techniques enable the execution of specific information processing tasks through a deep understanding of context and semantic relationships. In contrast to traditional recurrent neural networks and long short-term memory (LSTM) models, the architecture in this study mitigates performance degradation caused by long-term dependencies, enhancing both parallel computation and the capture of long-distance information. An innovation on this architecture, BERT, employs a bidirectional Transformer encoder that fully considers contextual information within the input.

In this study, BERT is leveraged to map each amino acid sequence into a feature vector, treating each molecular sequence in the reference dataset as a sentence and each amino acid as a word. The multi-head self-attention mechanism employed by BERT captures intricate relationships between every possible pair of amino acids, thereby enhancing feature representation. By utilizing transfer learning, BERT effectively applies insights from natural language processing to protein sequences, significantly reducing the reliance on extensive task-specific data while facilitating efficient feature extraction even in data-scarce scenarios. The selected pretraining model for this experiment is BERT-Small, consisting of 4 encoder layers, each with 8 attention heads and 512 hidden units. This process encompasses two primary steps: sequence embedding and feature extraction through the encoder layers. During the sequence embedding phase, [CLS] and [SEP] tokens are incorporated to ensure proper embedding within the BERT framework. As BERT processes each amino acid, it generates token, segment, and position embeddings, enabling the model to comprehend relationships among different amino acids in the sequence. Token embeddings represent words based on a specific tokenization methodology, while segment embeddings distinguish positions within the sequence. Position embeddings convey both relative and absolute positional information, as articulated in formulas (1) and (2), following the position vector framework proposed by Vaswani et al. (32).

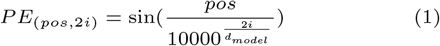

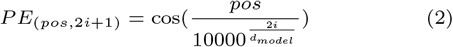

At the encoder layer stage, multi-head self-attention and feedforward neural networks are employed to process embedding vectors and transform them into more intricate feature representations. The multi-head self-attention sub-layer performs attention calculations at each position of every input sequence, capturing the relationships between the current position and all other positions.

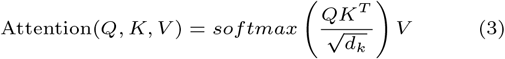

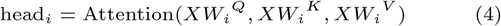

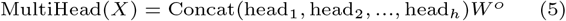

In the above formula, *Q, K*, and *V* , respectively, represent the matrices *Q*(*uery*), *K*(*ey*), and *V* (*alue*) obtained through different linear transformations of the input sequence. *W*_*i*_^*Q*^,*W*_*i*_^*K*^ ,*W*_*i*_^*V*^ and *W*_*i*_^*O*^ represent weight matrices, where *d*_*k*_ denotes the dimensionality of the key vectors in the *Key* matrix. The feedforward neural network (FFNN) sub-layer achieves information processing by applying nonlinear transformations to the contextual representations at each position. This sub-layer consists of an activation function, like *ReLU* , along with two linear transformations, facilitating the efficient extraction of features. The specific formula is as follows:

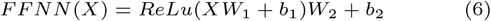

where *W*_1_ and *W*_2_ stand for weight parameters, *b*_1_ and *b*_2_ signify the bias terms. Additionally, *X* indicates the representation obtained after passing through the multi-head self-attention sub-layer.

### Network architecture post feature extraction

In this study, we constructed a function prediction network framework that integrates CNN, Bi-LSTM, attention mechanisms, and TCN. After protein sequences are encoded using BERT, the framework is employed to further enhance feature extraction. The CNN autonomously extracts feature information through one-dimensional convolutional layers, refining the data initially obtained from BERT. The outputs from the CNN are then directed to the Bi-LSTM layer for further processing.

The LSTM structure effectively captures long-range associations in protein sequences by incorporating memory cells and forget gate mechanisms. Memory cells retain essential information while the forget gates discard irrelevant features, thus improving the extraction of complex structures. Every Bi-LSTM unit accurately captures the temporal dependencies within the sequence by processing features both forward and backward. Given that protein sequences exert functional influences from both the N- and C-terminal ends, Bi-LSTM is particularly advantageous for tasks such as amino acid sequence prediction, as it can simultaneously extract critical information from both ends.

The attention mechanism enhances the model’s flexibility, allowing it to balance and selectively focus on different segments of the input when analyzing protein sequences. Key components include queries, keys, values, attention scores, softmax functions, and final attention outputs. The comparison between the query and key is computed, the results are normalized using the softmax function, and the normalized attention weights are applied to the values to produce the outputs that are ultimately required. In this context, BBATProt leverages self-attention to dynamically adjust the significance of each region in the sequence, thereby enhancing the model’s explainability. This approach clarifies how various parts of the protein sequence contribute to the final prediction, rendering the decision-making process more interpretable compared to conventional black-box models.

Following this, the flatten layer transforms the multidimensional input data generated by the attention mechanism into a one-dimensional array, which is then passed to the TCN layer, effectively connecting features across different layers and enhancing the model’s ability to represent the input data.

The core components of the TCN comprise one-dimensional convolutional layers, residual connections, dilated convolutions, and causal convolutions. The incorporation of residual and causal connections in TCNs prevents information leakage from future positions, maintaining a clear cause-and-effect relationship in sequence processing. The temporal relationships and positional relation yeilded from procession, reflecting the protein folding and conformational information, are critical for function . The model’s receptive field grows exponentially as the number of layers increases thanks to the dilated convolutions, facilitating efficient capture of global sequence dependencies. This design allows for a clearer trace of which segments of the sequence influence the output, thereby enhancing interpretability compared to other recurrent models.

The integration of Bi-LSTM and TCN in BBATProt is particularly advantageous because these architectures complement each other’s strengths in local and global feature extraction. While Bi-LSTM excels in capturing sequential and bidirectional dependencies, it can face challenges with parallelization and efficient processing of lengthy sequences. Conversely, TCN is adept at capturing long-range dependencies and enabling parallel processing, though it lacks the bidirectional insight provided by Bi-LSTM. By combining these two architectures, BBATProt effectively captures both local sequential information and long-range dependencies. This framework offers a more transparent perspective on how features from various segments of the sequence are processed and synthesized to produce the final prediction. This transparency is vital in biological applications, where comprehending the underlying mechanics takes precedence over merely achieving high predicted accuracy.

Ultimately, in the Dense layer, feature information from each layer is integrated, with dimensions progressively reduced in a hierarchical manner. The final classification output is achieved using a sigmoid activation function, ensuring that the features learned by the network are distilled into a concise result.

### Model Evaluation Parameters

Six metrics were taken into consideration in order to assess the performance of the classification models: F1 score (FSc), specificity (SPE), sensitivity (SEN), precision (PRE), accuracy (ACC), and Matthews correlation coefficient (MCC). These measures have the following definitions:

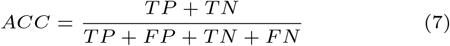

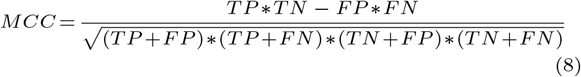

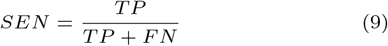

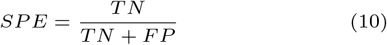

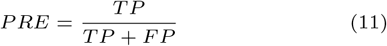

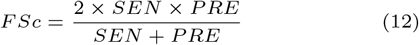

In the above formula(7)-(12), *TP* and *TN* designate the counts of correctly forecasted positive and negative instances respectively, while *FP* and *FN* denote the quantities of erroneously predicted positive and negative instances.

## Experimental Analysis

To comprehensively validate the universal applicability of the BBATProt framework in peptide and protein function prediction tasks, this study focused on three key categories: enzyme function prediction, peptide function prediction, and PTM site prediction. Five diverse datasets were utilized to ensure a robust evaluation of the model’s versatility, covering the prediction of various biological functions, including carboxyl ester hydrolytic activity, antimicrobial activity, dipeptidyl peptidase-lV inhibition, and potential sites for post-translational modifications such as lysine crotonylation (Kcr) and lysine glycation (Kgly).

### Comparison of BERT Representational Capability Across Different Sizes

Google Research has recently released pre-trained BERT models in multiple sizes (14). Meanwhile, Zhang et al. selected 556,603 protein data entries from UniProt to pretrain their model (33). In this study, through detailed comparative experiments, the goal is to determine the most suitable feature representation. Six representative models were selected for fine-tuning out of all the pre-trained BERT models: BERT-Tiny, BERT-Mini, BERT-Small, BERT-Base, and BERT-Protein. Specific parameters can be found in the supplementary data, and the comparative results are presented in the charts 2. Relatively, the Small-sized models show better results during training. Consequently, the pre-trained BERT-Small model was chosen to extract characteristics from protein sequence data.

### t-SNE Visualization of Features Extracted Layer by Layer by BBATProt on Protein Dataset

The core task of deep learning in sample classification is the continuous extraction of features with unique representations, a process that evolves throughout the entire network structure. To thoroughly depict the advancement of the BBATProt model in the domain of feature learning, the concept of t-distributed stochastic neighbor embedding (t-SNE) has been brought into light for the reduction of dimensionality and visualization of information prevalent in the samples (34). This particular pathway paves the way to a heightened understanding of the abstraction and learning of sample features across distinct network strata. To elucidate the hierarchical feature extraction within the network, the analysis progresses from the raw samples through the convolutional layer, Bi-LSTM layer, attention layer, and TCN layer, culminating in the final Dense layer.

In the visualization graph shown in Fig. 2, positive samples are represented by blue dots and negative samples by green dots. A significant observation from this experiment is that with each additional layer in the neural network structure, the deeper layers exhibit enhanced capabilities for feature extraction.

**Fig. 2.**
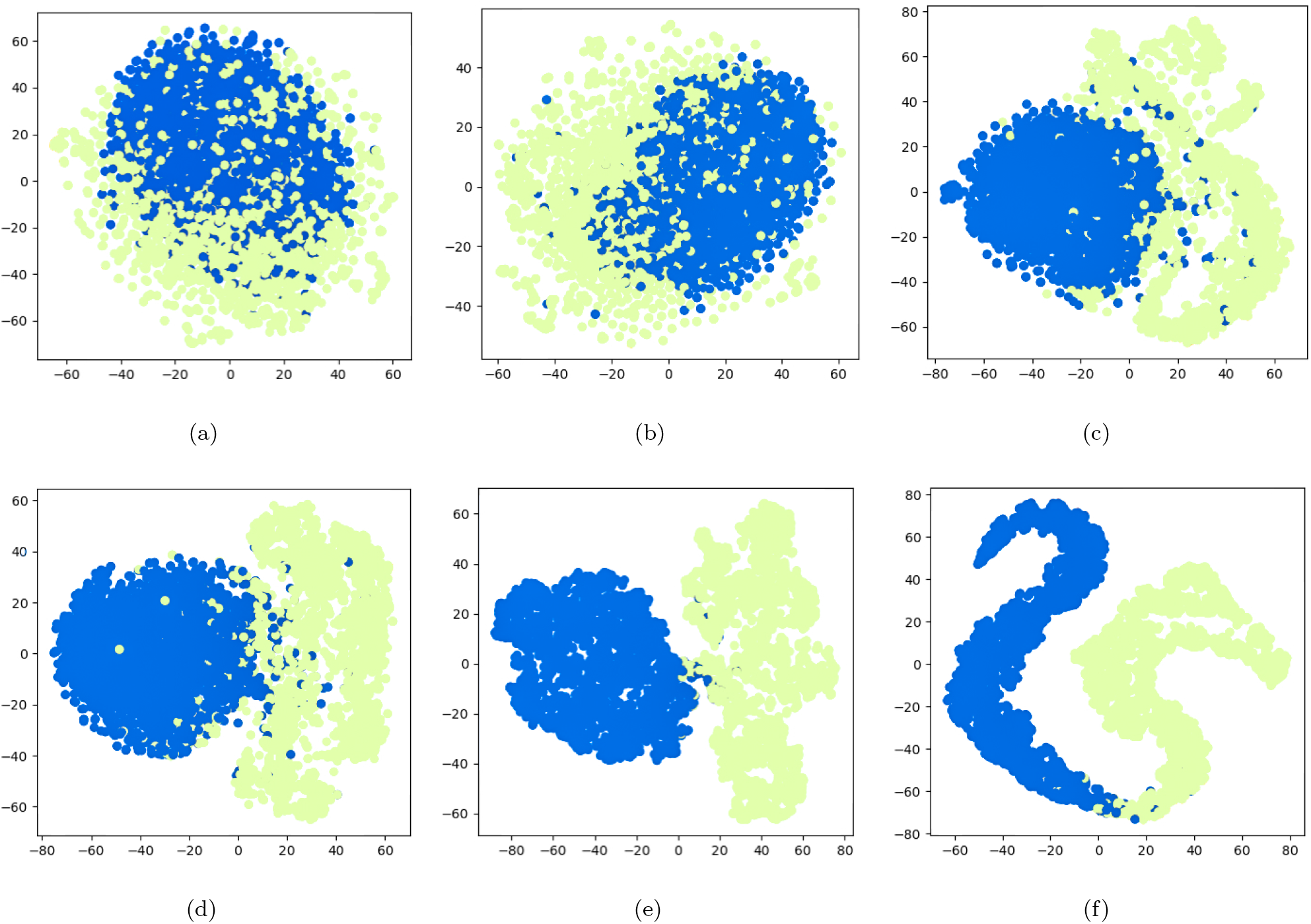
The t-SNE visualization of features extracted layer by layer through BBATProt on the protein dataset. (a) t-SNE visualization of encoded features after BERT encoding. (b) t-SNE visualization of features after the convolutional layer. (c) t-SNE visualization of features after the Bi-LSTM layer. (d) t-SNE visualization of features after the Attention layer. (e) t-SNE visualization of features after the TCN layer. (f) t-SNE visualization of features after the Dense layer.

**Fig. 3.**
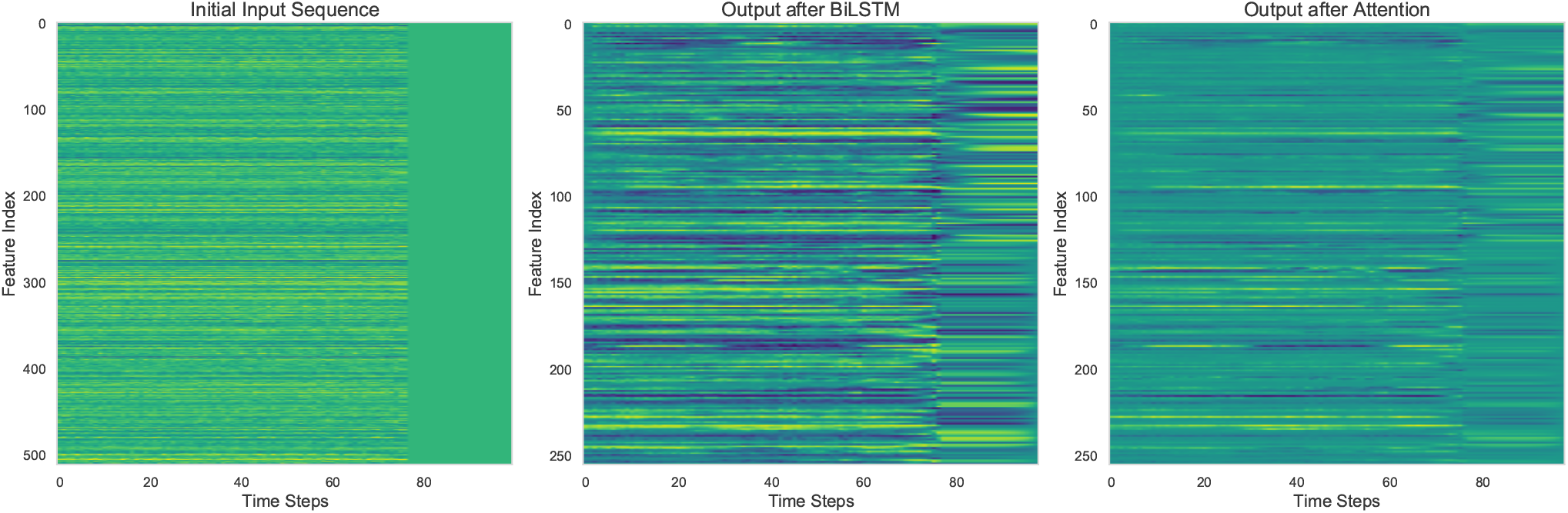
Visualization of the attention mechanism.

### Comparison of BBATProt with existing predictors on hydrolytic enzyme datasets

This segment starts with the formulation of a benchmark dataset containing carboxylesterases identified by EC number 3.1.1.1. Known for their ability to act as catalysts, these enzymes facilitate the hydrolysis of ester. Beyond their functions in biological processes like drug metabolism, food digestion, and xenobiotic detoxification, these enzymes play a crucial role in various industries, including renewable energy, food processing, and bioremediation in environmental engineering. Given the extensive time and effort required by traditional experimental methods, computational approaches offer valuable and efficient alternatives.

The positive dataset was obtained from the UniProt. In order to improve feature extraction, sequences exceeding a length of 512 were excluded, focusing on those within the range of 0-512. Furthermore, to reduce the influence of high sequence similarity on the experimental results, CD-HIT was utilized to filter out redundant sequences with a similarity threshold above 80%. Negative samples were randomly generated from Swiss-Prot, maintaining the same length range as positive samples. These negative samples also underwent CD-HIT processing with a threshold set at 30%. Ultimately, a training dataset comprising 6960 hydrolytic enzyme samples was obtained, with 3480 each for positive and negative samples.

To demonstrate the effectiveness of the proposed network model, a comparison was drawn against various conventional machine learning classification strategies, including KNN, RF, SVM, and XGBoost. Hydrolytic enzymes were selected specifically for a 10-fold cross-validation, the comprehensive results of which are presented in Table 3. As these data reveal, BBATProt excels in the intricate domain of enzyme protein function prediction, exhibiting peak performance across a range of evaluation standards.

**Table 3.** 10-Fold Cross-Validation Comparison Between BBATProt and Other Machine Learning Methods on the Hydrolysis Enzyme Dataset with the Same BERT Encoding Features(%).

| Predictor | ACC | MCC | SEN | SPE | PRE | FSc | AUROC |
| --- | --- | --- | --- | --- | --- | --- | --- |
| BBATProt | **96.60** | **93.33** | **95.59** | **97.79** | **97.97** | **96.71** | 96.63 |
| BERT+RF | 75.57 | 51.30 | 71.50 | 79.65 | 77.87 | 74.53 | 97.97 |
| BERT+XGBOOST | 80.63 | 61.28 | 79.25 | 82.02 | 81.51 | 80.34 | **99.58** |
| BERT+KNN | 79.35 | 59.87 | 88.92 | 69.88 | 74.66 | 81.13 | 86.35 |
| BERT+SVM | 83.79 | 67.61 | 82.38 | 85.23 | 84.78 | 83.54 | 84.60 |

Note: The optimal performance for each metric is indicated in bold.

Additionally, a comparative analysis was conducted to evaluate the encoding effectiveness of two NLP models, BERT and Word2Vec (35), on the hydrolase dataset. Both models were applied to encode the data, and their performance was assessed using an identical neural network architecture for classification. Word2Vec uses a dual-modality that encompasses both skip-gram and CBOW strategies. Continuous Bag of Words (CBOW) mainly concentrates on forecasting the desired term when provided with contextual clues, whereas the skip-gram tactic emphasizes the estimation of the surrounding wording based on the specific target term. Table 4 shows the efficacy measures of each system via a 10-fold cross-validation.

**Table 4.** Comparative Results of 10-Fold Cross-Validation on the Hydrolysis Enzyme Dataset Using Different NLP-based Feature Representation Models(%).

| Predictor | ACC | MCC | SEN | SPE | PRE | FSc | AUROC |
| --- | --- | --- | --- | --- | --- | --- | --- |
| BBATProt | **96.60** | **93.33** | **95.59** | **97.79** | **97.97** | **96.71** | **96.63** |
| Word2Vec-CBOW+Conv+BiLSTM+Att+TCN+Dense | 48.36 | NA | 50.00 | 50.00 | 24.18 | 32.59 | 48.36 |
| Word2Vec-Skip-gram+Conv+BiLSTM+Att+TCN+Dense | 48.22 | NA | 60.00 | 40.00 | 29.11 | 39.19 | 48.22 |

Note: The optimal performance for each metric is indicated in bold. NA is used as a representation when the result is not available.

The outcome decisively attests that the pre-trained BERT model is superior to the other two NLP models in every performance measurement, strongly underscoring the notable effectiveness of BERT encoding compared with Word2Vec.

### Comparison of BBATProt with existing predictors on peptide datasets

The assessment of BBATProt’s predictive capability on peptide datasets necessitates a comprehensive examination of both the AMP and DPP-IV inhibitory peptide datasets.

#### Comparison of BBATProt with existing predictors on antimicrobial peptide datasets

In this section, we provide a detailed comparison of BBATProt with 13 SOTA methods (36; 37; 38; 39; 40; 41; 42; 43; 44; 45; 26) using the XUAMP dataset. This comparison was conducted through a comprehensive evaluation in an independent testing environment. The results demonstrate that BBATProt significantly outperforms the other methods across key metrics, including ACC, MCC, SEN, Fsc, and the AUROC. As shown in Table 5, BBATProt shows an improvement ranging from 2.81% to 31.96% in ACC, 5.42% to 63.68% in MCC, 0.56% to 64.4% in SEN, 2.39% to 49.46% in FSc, and 0.71% to 40.51% in AUROC when compared to its counterparts.

**Table 5.**
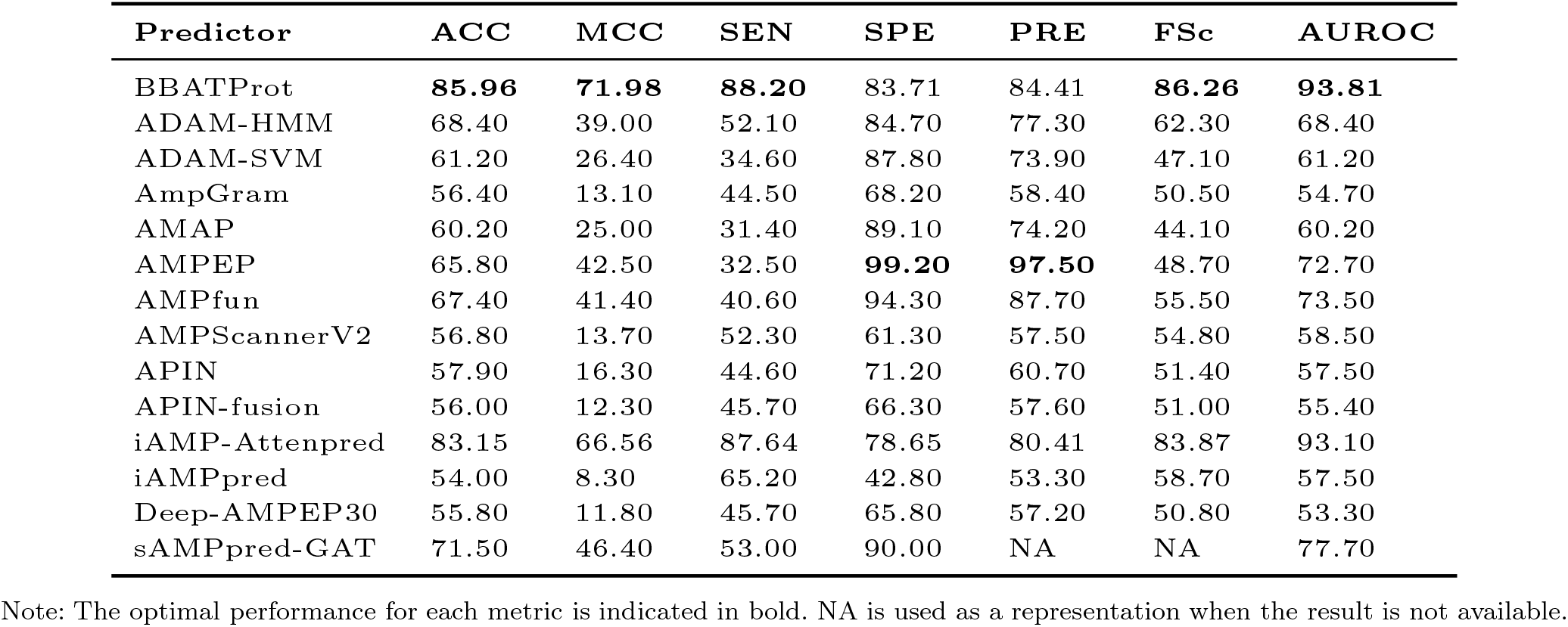
Performance Comparison Results of BBATProt and Existing Methods on Independent AMP Dataset(%).

Additionally, BBATProt ranked 6th and 2nd in the SPE and PRE evaluations, respectively. While AMPEP achieved the highest scores in these two metrics, its overall average performance across the other five evaluation metrics was 32.80% lower than that of BBATProt. This indicates that BBATProt provides a more comprehensive extraction of feature information, emphasizing its strengths across various predictive tasks and suggesting potential avenues for further improvement in protein analysis. By examining the model characteristics of AMPEP in more detail, valuable insights can be gained for optimizing BBATProt in future studies. Moreover, the visualization of the attention mechanism, as shown in Figure 4, reveals that BBATProt effectively filters out irrelevant information during the computation process, demonstrating a focused and refined approach to data analysis. This visual evidence supports the conclusion that the attention mechanism is crucial for enhancing BBATProt’s ability to accurately predict protein functions by concentrating on the most relevant features.

**Fig. 4.**
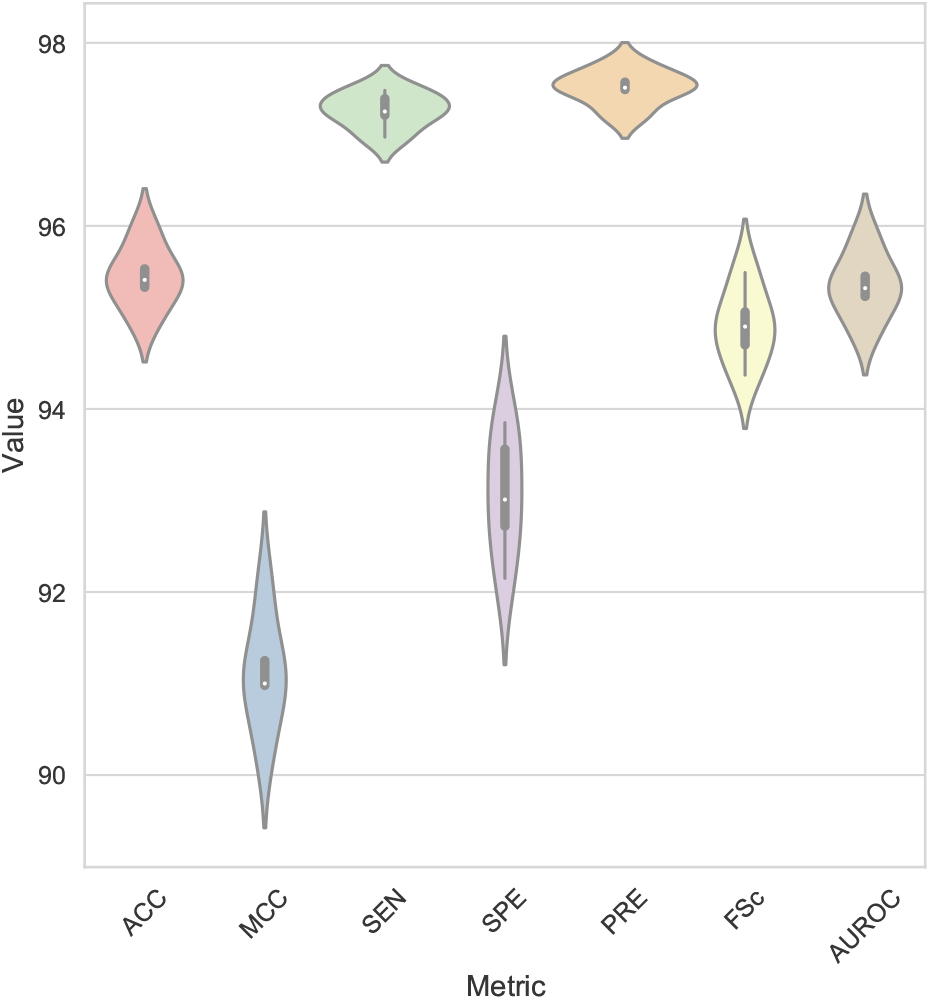
Five times and average performance results of 10-fold cross validation method based on the AMP benchmark dataset(%).

**Fig. 5.**
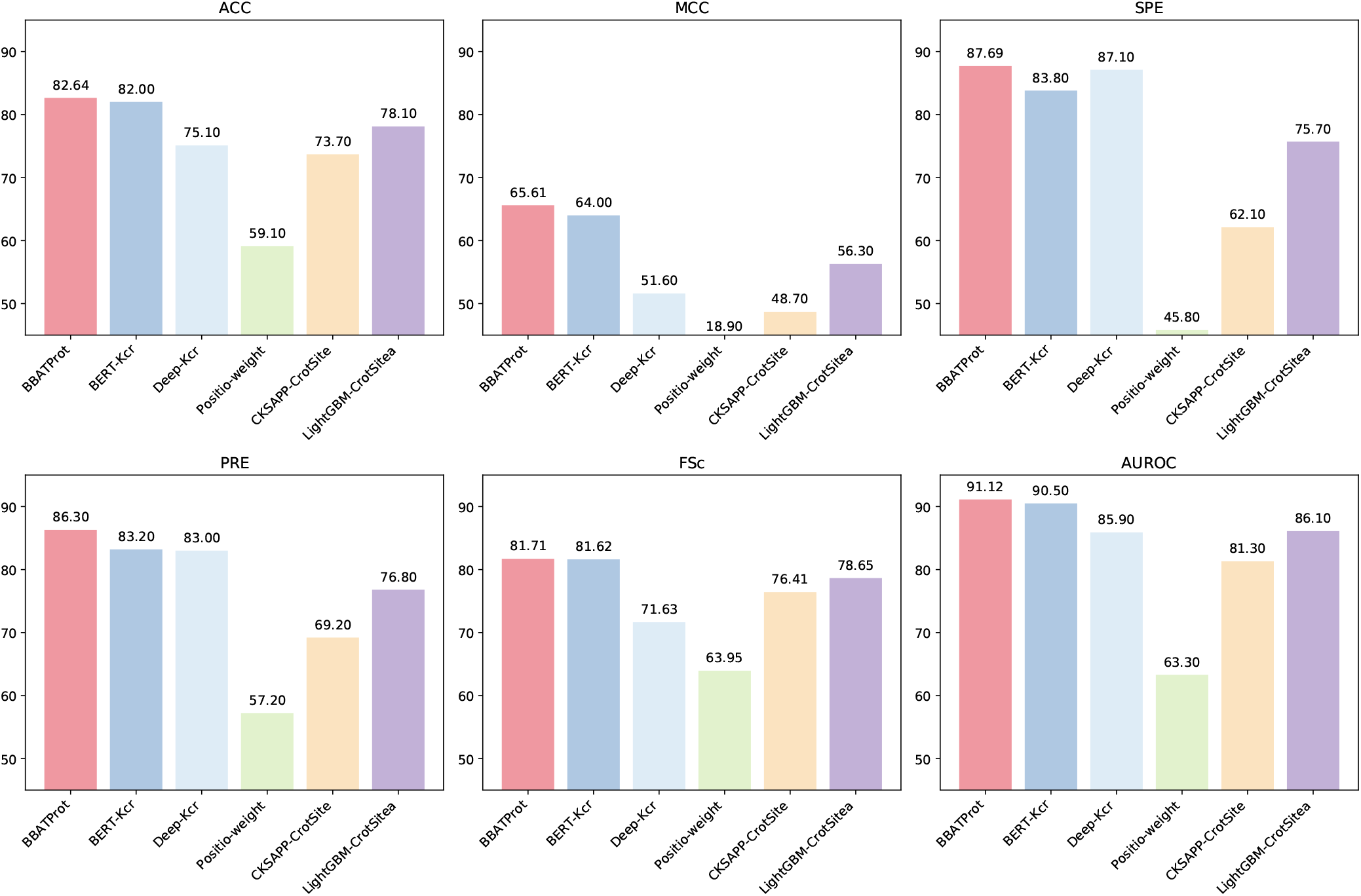
Performance Comparison Results of BBATProt and Existing Methods on Independent Kcr Site Prediction Dataset(%).

To comprehensively assess the effectiveness of BBATProt in AMP prediction, a benchmark dataset was constructed. During dataset construction, sequences with more than 20 natural standard amino acids were excluded to ensure data consistency. The AMP dataset was compiled from various sources, including the AMPer, APD3, and ADAM databases (35; 46; 47). Data cleaning involved setting a CD-Hit threshold of 90% to remove redundant sequence information. Correspondingly, non-AMP data were sourced from UniProt, with a constraint on the residue length of protein fragments between 5 and 100 to maintain sequence lengths similar to those of the AMP dataset for experimental validity. The CD-Hit redundancy removal threshold for the non-AMP dataset was set to 40%. In addition, sequences with annotations containing terms such as ‘Defensin‘, ‘Antimicrobial‘, ‘Antibiotic‘ or ‘Antifungal‘ were excluded to ensure data purity.

Using the benchmark dataset, multiple repetitions of 10-fold cross-validation were performed to accurately evaluate the performance and generalization capability of the predictor. The resulting averages for each validation and across five repetitions are detailed in Figure . This method ensures a robust assessment of the performance of the model on various dataset subsets, emphasizing the reliability and repeatability of the results.

#### Comparison of BBATProt with existing predictors on inhibitory peptide datasets

Due to their crucial role in pharmaceutical development and diabetes treatment, distinguishing between inhibitory and non-inhibitory peptides of dipeptidyl peptidase-IV (DPP-IV) has become a key focus. The BBATProt model was trained and evaluated on a dataset designed to differentiate these two classes, including both inhibitory DPP-IV peptides and non-inhibitory counterparts (27). The dataset was split into a benchmark set and an independent test set, each containing an equal distribution of both peptide types. Specifically, the benchmark set comprised 532 samples, while the independent set contained 133 samples. As presented in Table 6, BBATProt outperforms other models in terms of ACC, MCC, SEN, and SPE, and shows comparable AUROC performance to the current SOTA models.

**Table 6.**
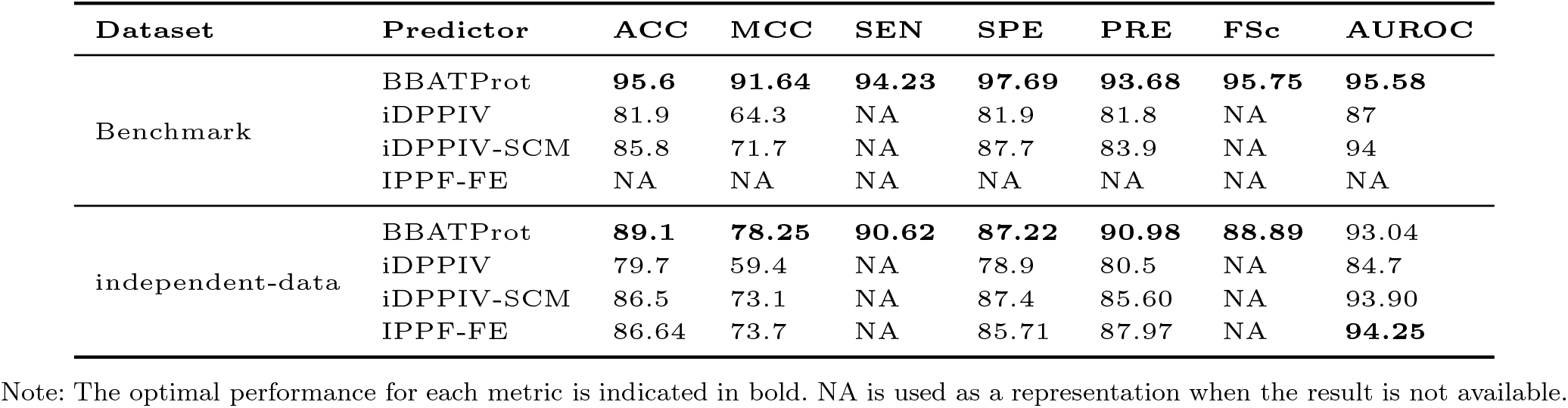
Performance Comparison Results of BBATProt and Existing Methods on the DPP-IV Inhibitory Peptide Dataset(%).

| Dataset | Predictor | ACC | MCC | SEN | SPE | PRE | FSc | AUROC |
| --- | --- | --- | --- | --- | --- | --- | --- | --- |
| Benchmark | BBATProt | **95.6** | **91.64** | **94.23** | **97.69** | **93.68** | **95.75** | **95.58** |
|  | iDPPIV | 81.9 | 64.3 | NA | 81.9 | 81.8 | NA | 87 |
|  | iDPPIV-SCM | 85.8 | 71.7 | NA | 87.7 | 83.9 | NA | 94 |
|  | IPPF-FE | NA | NA | NA | NA | NA | NA | NA |
| independent-data | BBATProt | **89.1** | **78.25** | **90.62** | **87.22** | **90.98** | **88.89** | 93.04 |
|  | iDPPIV | 79.7 | 59.4 | NA | 78.9 | 80.5 | NA | 84.7 |
|  | iDPPIV-SCM | 86.5 | 73.1 | NA | 87.4 | 85.60 | NA | 93.90 |
|  | IPPF-FE | 86.64 | 73.7 | NA | 85.71 | 87.97 | NA | **94.25** |

Note: The optimal performance for each metric is indicated in bold. NA is used as a representation when the result is not available.

### Comparison of BBATProt with existing predictors on PTM site prediction datasets

Concerning PTM prediction challenges such as protein lysine crotonylation and protein lysine glycation, these alterations can affect the functionality, stability, and affinity of proteins, subsequently steering their biological operations. However, experimental methods for site prediction are both expensive and time-consuming. In contrast, computational methods can provide reasonable predictions in a highly efficient and cost-effective manner. Using bioinformatics tools and algorithms, protein sequences and structures can be analyzed to predict potential modification sites.

To validate the advanced nature of the designed network structure, BBATProt was compared with some classical machine learning classification methods, including KNN, RF, SVM, and XGBoost. A 10-fold cross-validation was implemented for Kcr site prediction, with the outcomes detailed in Table 7. The evidence indicates the superior effectiveness of BBATProt, most notably in brief site prediction duties. This can be attributed to the fact that BBATProt is based on hierarchical learning, which allows it to adapt to protein data of different lengths and extract high-level features, making the model more effective in handling complex classification tasks.

**Table 7.** 10-Fold Cross-Validation Comparison Between BBATProt and Other Machine Learning Methods on the Kcr Site Prediction Dataset with Consistent BERT Encoding Features(%).

| Predictor | ACC | MCC | SEN | SPE | PRE | FSc | AUROC |
| --- | --- | --- | --- | --- | --- | --- | --- |
| BBATProt | **94.95** | **89.93** | **94.05** | **95.86** | **95.80** | **94.91** | 94.95 |
| BERT+RF | 74.34 | 49.49 | 83.36 | 65.34 | 70.63 | 76.46 | 96.01 |
| BERT+XGBOOST | 76.86 | 54.10 | 82.80 | 70.91 | 74.00 | 78.15 | **97.38** |
| BERT+KNN | 70.12 | 40.23 | 69.48 | 70.74 | 70.36 | 69.91 | 78.70 |
| BERT+SVM | 79.47 | 59.30 | 84.97 | 73.97 | 76.56 | 80.53 | 92.40 |

Note: The optimal performance for each metric is indicated in bold.

To avoid overestimating the performance of BBATProt, a comparison was made between its training results and those of other existing SOTA models on independent test sets for both Kcr and Kgly site prediction problems. In Table 5, BBATProt demonstrates superior performance across almost all metrics compared with the other models specifically designed for Kcr site prediction (28; 50; 51). Although it lags slightly behind the CKSAPP CrotSite algorithm in terms of SEN, this may be attributed to the lack of specific parameter tuning in BBATProt for addressing Kcr site prediction issues. This conclusion provides a potential avenue for future improvement research. Notably, in terms of ACC, MCC, SPE, PRE, FSc, and AUROC, BBATProt exhibits improvements of 8.94%, 16.91%, 25.56%, 5.3%, and 9.82%, respectively, compared with the CKSAPP CrotSite algorithm model. This underscores the significant overall performance advantage of BBATProt.

In the Kgly site prediction task presented in Figure 6, BBATProt demonstrates a clear superiority over other advanced SOTA predictors across all evaluated metrics (29; 52; 20; 53). Notably, for the Sensitivity (SEN) metric, BBATProt shows an improvement range of 18.9% to 56.9% compared to other predictors, underscoring its robust capability to accurately capture positive samples. This performance highlights the significant feasibility of BBATProt in post-translational modification (PTM) site prediction, emphasizing its effectiveness in identifying relevant sites with high accuracy.

**Fig. 6.**
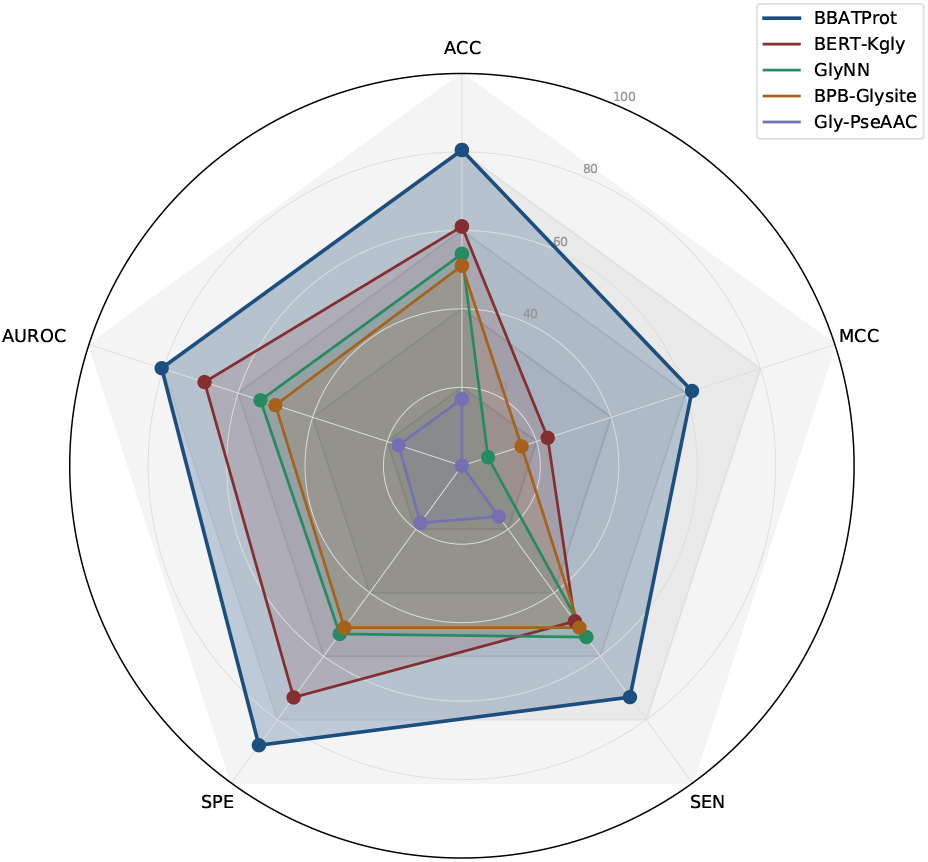
Performance Comparison Results of BBATProt and Existing Methods on the Kgly Site Prediction Dataset.(%).

## Conclusion

The function prediction of protein and peptide based on amino acid sequences is crucial in both academic research and industrial applications. This study introduces BBATProt, an innovative framework designed to tackle this challenge. Diverse biological datasets were transformed into peptide and protein datasets with varying amino acid sequence lengths, targeting prediction tasks such as hydrolases, AMP, DPP-IV inhibitory peptides, and PTM at Kcr and Kgly sites.

Comprehensive evaluations across multiple datasets confirm its superior accuracy, robustness, and generalization capabilities compared to SOTA models. Furthermore, this study enhances the interpretability of what is typically a black-box model by leveraging visualization techniques. The sequential network architecture, built around the spatial conformations of proteins, further validates the model’s effectiveness. Additionally, enhanced feature extraction through transfer learning has proven instrumental in improving prediction performance. Together, these approaches ensure that BBATProt not only excels in predictive power but also provides a more explainable and transparent model, offering deeper insights into how specific features of amino acid sequences contribute to functional predictions.

Although BBATProt performs well in the function prediction of protein and peptide, there is still room for improvement. Future research will focus on feature integration, further optimizing prediction performance by incorporating structural information of peptides or proteins into the integrated model’s input. Additionally, plans include the development of an intuitive and user-friendly web server to offer public prediction services, enhancing accessibility to a wider researcher base and expanding practical applications.

## Acknowledgments

This work was supported in part by the National Science Fund for Distinguished Young Scholars of China under Grant 62225303; in part by the National Natural Science Funds of China under Grant 62303039, 32371325 and 62433004; in part by the China postdoctoral Science Foundation BX20230034 and 2023M730190; in part by the Fundamental Research Funds for the Central Universities buctrc202201, QNTD2023-01; in part by Beijing Natural Science Foundation (L241014) and in part by the High Performance Computing Platform, College of Information Science and Technology, Beijing University of Chemical Technology.

## Data Availability

Publicly available datasets were analyzed in this study. These data can be found here: https://github.com/Xukai-YE/BBATProt.

## Key Ponits

- BBATProt proposes an explainable neural network framework for protein function prediction.
- The model outperforms state-of-the-art methods in various protein and peptide prediction tasks.
- Visualization confirms model interpretability, highlighting attentions focus on relevant data.

